# Three-dimensional imaging and quantification of mouse ovarian follicles via optical coherence tomography

**DOI:** 10.1101/2023.01.23.525192

**Authors:** Marcello Magri Amaral, Aixia Sun, Yilin Li, Chao Ren, Anh Blue Truong, Saumya Nigam, Zexu Jiao, Ping Wang, Chao Zhou

## Abstract

Ovarian tissue cryopreservation has been successfully applied worldwide for fertility preservation. Correctly selecting the ovarian tissue with high follicle loading for freezing and reimplantation increases the likelihood of restoring ovarian function, but it is a challenging process. In this work, we explore the use of three-dimensional spectral-domain optical coherence tomography (SD-OCT) to identify different follicular stages, especially primary follicles, compare the identifications with H&E images, and measure the size and age-related follicular density distribution differences in mice ovaries. We use the thickness of the layers of granulosa cells to differentiate primordial and primary follicles from secondary follicles. The measured dimensions and age-related follicular distribution agree well with histological images and physiological aging. Finally, we apply attenuation coefficient map analyses to significantly improve the image contrast and the contrast-to-noise ratio (p < 0.001), facilitating follicle identification and quantification. We conclude that SD-OCT is a promising method to noninvasively evaluate ovarian follicles.

## 1. Introduction

Worldwide, women have an 18.55 % lifetime risk of cancer, and breast cancer is responsible for 11.7 % of all cases^1^. About 10 % of cancers occur in women younger than 45 years old^2^, their most fertile years. Malignant cancer is mainly treated with chemotherapy and radiotherapy. Despite advances in these therapies and increasing survival prognoses, depending on the woman’s follicular reserve, age, and the drugs used, cancer still poses a risk of premature ovarian insufficiency (POI), reducing fertility. Additionally, some conditions, such as autoimmune and hematological diseases that require chemotherapy or radiotherapy, bone marrow transplantation, recurrent endometriosis, family history of premature menopause, or even social choices (like wishing to postpone pregnancy) also increase the risk of POI^2,3^. Preserving fertility is important to these patients, and thus options for fertility preservation have attracted growing interest^4^ .

Ovarian tissue cryopreservation (OTC) is one promising technology for fertility preservation. Briefly, OTC involves collecting ovarian tissue and preserving it in liquid nitrogen while the patient receives their cancer treatment. When the treatment is finished, the ovarian tissue is defrosted and reimplanted, where it can produce new eggs and eventually lead to a normal conception. OTC allows the storage of a large number of primordial and primary follicles. It can be performed at any time of the menstrual cycle and has been proposed mainly for prepubertal females^3^. OTC has also been recommended for patients who cannot delay cancer treatment long enough to undergo ovarian stimulation and oocyte retrieval^3^. OTC has been successful even after several years of tissue storage, and its use as an alternative (or complement) to oocyte/embryo cryopreservation has developed rapidly over the last two decades ^5^ .

While OTC is advancing technically, optimizing it remains challenging. Cryopreservation of ovarian tissue theoretically can efficiently preserve thousands of ovarian follicles at one time. However, follicular densities and stages are not evenly distributed and depend on the ovary site. Currently, the tissue site to be preserved and later reimplanted is selected under a stereomicroscope. Although this technique for localizing and quantifying follicles within the tissue is feasible, the dense and fibrous nature of ovarian tissue makes it labor-intensive and unreliable^6^. Therefore, a method to identify the presence and density of follicles before cryo-storage and after thawing could ensure that only tissue with a high follicle loading was utilized. Such a method would also help further optimize clinical cryopreservation protocols. At the same time, any *in situ* follicle identification method must not harm tissue or compromise long-term follicle survival and growth.

Optical coherence tomography (OCT) is a promising candidate for identifying follicle presence and quantifying follicle density. Without the use of contrast agents, it can non-invasively provide three-dimensional (3D) images with micrometer scale resolution, yielding millimeter-scale depth information. It has been largely adopted in the ophthalmic field for imaging retina-related disease^7,8^, but also has found applications in other fields, such as dermatology^9^, rheumatology^10^, dentistry^11^, cardiovascular medicine^12^, and in developmental biology and organoid models^13^. OCT may be employed in selecting ovary tissue with the highest density of primordial follicles for cryopreservation and/or reimplantation. Future OTC applications may benefit from the non-invasive and 3D imaging capability of OCT to distinguish different follicle stages and densities.

As in humans, advanced maternal age is associated with a decrease in ovarian function incidence in naturally aged mice. Using a natural reproductive aging mouse model, this work explores OCT’s ability to quantitatively measure and select ovary tissue by characterizing different follicle stages and differences in follicular distribution. We use 3D spectral-domain OCT (SD-OCT) to identify different follicle stages, compare them to H&E images, and measure the size and age-related density distribution differences in mouse ovaries. The granulosa cell layer thickness is used to differentiate between primordial/primary and secondary stages. Finally, we apply attenuation coefficient map analyses to improve the image contrast and facilitate follicle identification from 3D OCT images.

## 2. Methods

### Mouse ovary sample preparation

Twelve female NOD/SCID mice (Jackson Laboratory, Bar Harbor, ME), divided into four age groups (18-day-old, 4, 12, and 23-week-old; n = 3 per age group) were housed in a 12-hour light/dark cycle at 22 °C. All animal experiments were performed in compliance with guidelines approved by the Institutional Animal Care and Use Committee (IACUC) at Michigan State University. Paired ovaries were harvested and were fixed in 4 % paraformaldehyde overnight, and kept in 70 % ethanol at 4 °C.

### OCT imaging system

The ovary samples were imaged using a custom-made SD-OCT system. The system consisted of a superluminescent light diode (SLD) (cBLMD, Superlum - Cork, Ireland), operating at an 850 nm central wavelength with a 160 nm bandwidth, producing 15 mW of optical power. The output light was coupled to a 50:50 fiber coupler (Hi780, AC Photonics, Inc – CA) that split the light between the reference and sample arms of the system. The reference arm consisted of a fiber collimator (F220APC-850, Thorlabs, Inc – NJ) to collimate the light from the fiber output, a variable neutral density (ND) filter, a 50 mm achromatic lens (AC254-050-B-ML, Thorlabs, Inc – NJ) to focus the light onto the reference mirror, and dispersion compensation glass. The sample arm consisted of a fiber collimator (F220APC-850, Thorlabs, Inc – NJ), a pair of galvanometer scanners (GVS002, Thorlabs, Inc – NJ), a 100 mm lens (AC508-100-B-ML, Thorlabs, Inc – NJ), a 75 mm lens (AC508-075-B-ML, Thorlabs, Inc – NJ), and a 5x (M Plan Apo NIR 5x, Mitutoyo - Japan) or 10x objective lens (M Plan Apo NIR 10x, Mitutoyo - Japan). The backscattered light from the sample and the reflected light from the reference arm interfered and were detected by a spectrometer (Cobra-S 800 OCT Spectrometers - Wasatch Photonics, Logan, UT) and a 2048-pixel, 80 kHz USB line-scan camera (Teledyne e2V - OctoPlus). Custom-made software controlled and synchronized the acquisition and data processing. This system provided a lateral resolution of ∼3.48 µm, an axial resolution of ∼2.48 µm in air, and ∼93 dB sensitivity.

### Image acquisition

3D OCT datasets of all samples were acquired with a FOV of ∼3.3 mm × 3.3 mm, using 2000 A-scans per B-scan and 2000 B-scans, and 1.8 mm in depth with 1014 pixels. For whole ovary 3D reconstruction, all mouse ovary samples were measured while immersed in 1X PBS to avoid dehydration and reduce the reflectance of the first air-tissue interface, improving the image quality. Stereomicroscope images were acquired in the same orientation as the OCT images. The sample was marked with blue ink to guide histology alignment and posterior image registration and comparison.

### OCT Image processing

The OCT 3D datasets were filtered with a median filter (3×3 window), to reduce speckle noise, and resliced to *en face* view (top view), to match the histology viewing orientation. A moving average of every five depth positions was calculated to improve the contrast of the ovary structure, allowing better OCT and histology image comparison. The histology and OCT images were aligned to identify the different stages of follicles, and representative images are presented and discussed.

### Histology

For histology, tissues excised from healthy ovaries were paraffin-embedded, sectioned (5 µm thick), and stained conventionally with hematoxylin and eosin (H&E). The sections were scanned by an Aperio VERSA (Leica Biosystems, Deer Park, IL), a digital pathology whole tissue–slide scanner. Images were analyzed using Aperio ImageScope software (Leica Biosystems, Deer Park, IL). This software uniquely allows the VERSA imaging system to navigate, annotate, and analyze any field or all fields of the whole tissue section within its original context in a single image file.

### Attenuation Coefficient Map (ACM)

OCT image contrast relies on the intensity of the light backscattered by biological tissue. Thus, two effects can decrease the follicle’s visibility. The first effect is due to the OCT signal intensity exponential decay as light travels through the biological tissue (Beer’s law). As a result, structures at the same depth with similar scattering potentials have similar intensities, but structures with similar scattering potentials at different depths present different intensities, resulting in low contrast among these structures. The second effect is due to highly attenuating structures (such as blood vessels or high cellular concentration) that reduce the amount of light that can penetrate deeper into the tissue. This is a local effect that can shadow structure and reduce the contrast of shadowed follicles. One approach to correcting these artifacts and improving the contrast is to obtain the attenuation coefficient map (ACM), which changes the image contrast source from the backscattering intensity to the attenuation coefficient of the tissue for every image pixel. The attenuation coefficient is a property of the biological tissue itself, and it is independent of the OCT setup or light power level. It estimates the decrease in light intensity as it travels through biological tissue. In this work, we employed the ACM ^14–16^ to correct image artifacts and enhance the contrast of follicle structures. After applying an ACM, the attenuation and shadowing effects were corrected. Furthermore, we measured the contrast and contrast-to-noise ratio (CNR) of the granulosa layer of 80 PF in different depths to evaluate the image quality improvement by comparing the ACMs with structural OCT images.

### Statistical Analysis

The follicle’s dimension measurements were performed using Fiji image processing software. All the group comparisons were performed by paired t-test. All the values are presented as mean ± standard deviation (STD). All data in graphs are presented as mean ± STD. The statistical analysis was done with MATLAB 2022b (The MathWorks, Inc., Massachusetts, USA).

## 3. Results

### 3D OCT imaging of mouse ovaries

Figure 1a presents OCT *en face* projections, stereomicroscope images, and OCT 3D rendered images of representative mouse ovaries for each age group (18-day-old, 4, 12, and 23-week-old). As expected, the stereomicroscope image (Figure 1a central column) does not provide much information regarding the follicle structures inside the mouse ovaries. The OCT 3D renderings (Figure 1a right column) provide more information about the morphology of the ovary, making it possible to visualize the follicles at different depths. The OCT projections (Figure 1a left column) collapse the 3D information into a single 2D image, losing the depth dimension. However, they highlight the follicle structures inside the ovary tissue (circular structures indicated by red arrows in Figure 1) in a single image, which will be useful in speeding tissue selection in future clinical applications. Figure 1b presents a representative orthogonal view from the 3D OCT dataset (the 4-week-old mouse dataset presented in Figure 1a), demonstrating OCT’s ability to recover the 3D morphology of the ovarian tissue. The 3D rendering of this data set can be found in the Supplementary Video.

**Figure 1:**
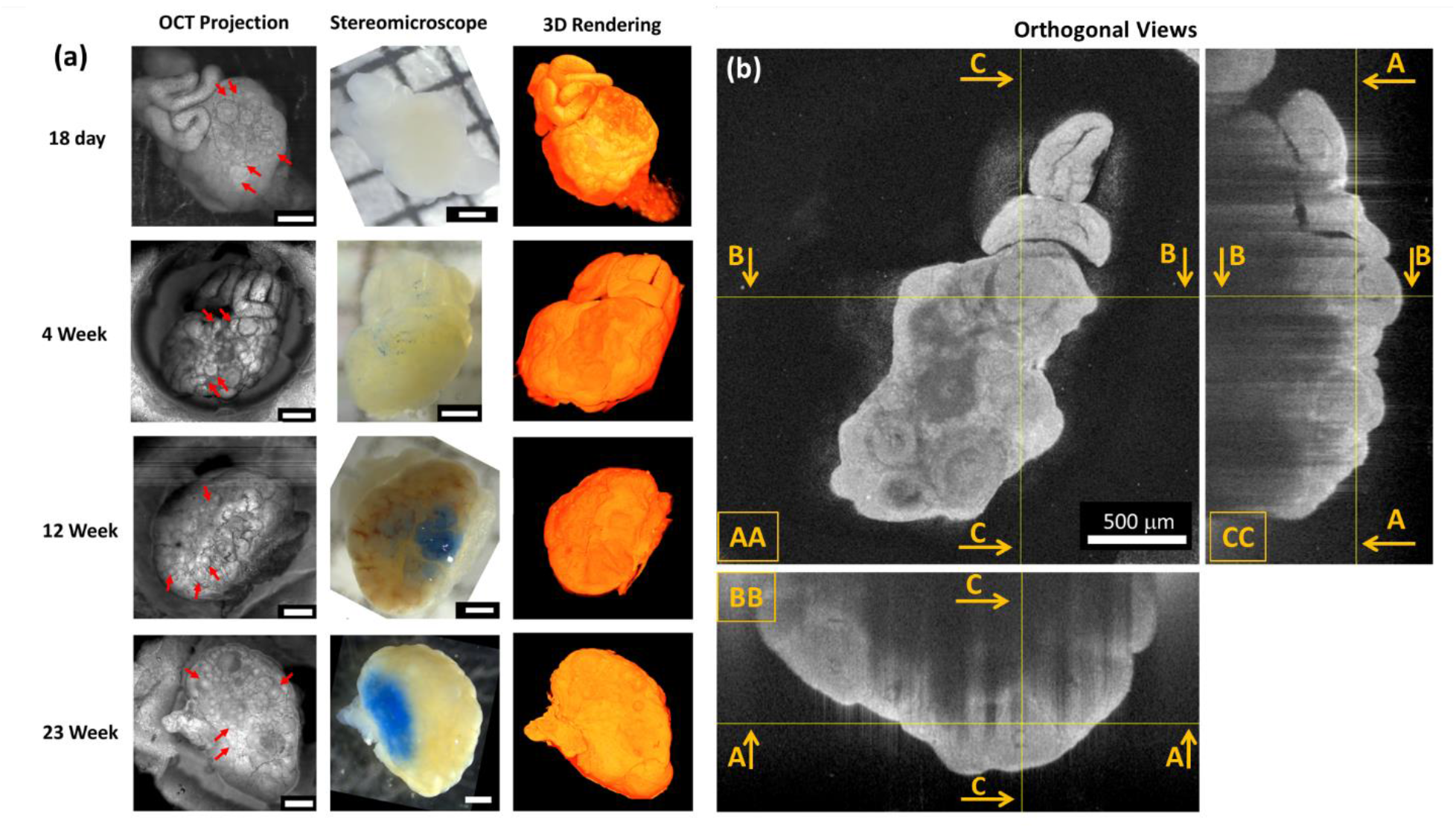
(a) En face projections of an OCT image (left) and stereomicroscope images (center) and OCT renderings (right) of 18-day-old, 4, 12, and 23-week-old mice ovaries. Red arrows point to examples of follicles highlighted by the OCT projection. (b) Three orthogonal views of the 3D OCT dataset: ([AA] top view, [BB] front view, [CC] left view. Scale bar, 500 µm.

### Correlation of OCT images with H&E staining

Figure 2 presents a side-by-side comparison of the H&E histology and the obtained OCT *en face* view of registered regions, from which it is possible to identify the three stages of follicle development inside the mouse ovary tissue. The yellow and blue arrows in Figure 2 a-d point to primary (PF) and secondary follicles (SF), respectively. The PF and SF are very similar, differing mainly in size and the number of their cellular layers. The PF has one layer of granulosa cells, as shown in the H&E histology. SF are larger, have two or more layers of granulosa cells, and may present small accumulations of fluid in the intracellular spaces (follicular fluid). The follicular fluid in OCT appears as black circles/rings, and the granulosa cells appear as bright circles/rings in the follicle region (Figure 2c and d). The green arrow in Figure 2 e and f points to the antral follicles (AF), characterized by the presence of an antrum (fluid-filled cavity), which is the blank region inside the follicle in the H&E image (Figure 2e) and the corresponding darker region in the OCT images(Figure 2f). The oocyte inside the AF can also be easily identified in the OCT images (red arrow).

**Figure 2:**
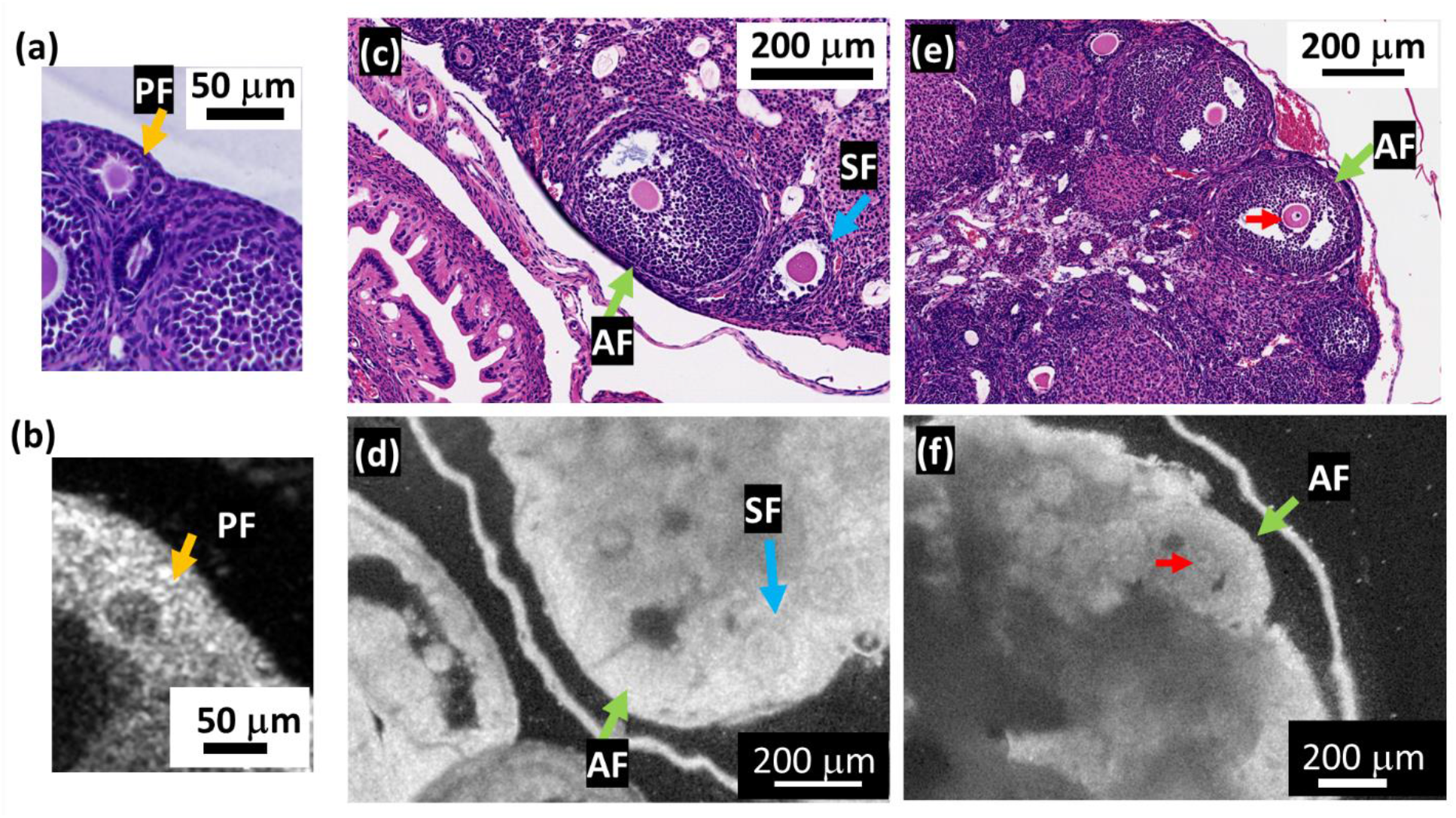
Comparative H&E histological image (a,c, and e) and OCT en face images (b,d, and f). PF, primary follicle (yellow arrow); SF, secondary follicle (blue arrow); AF, antral follicle (green arrow); oocyte (red arrow).

### OCT characterization of ovarian tissue from four different aged groups

The difference in the OCT and H&E images from the four age groups is presented in Figure 3. For 18-day-old, 4, 12, and 23-week-old ovarian tissues, (a), (b), (c), and (d) are representative H&E histological images, and (e), (f), (g), and (h) are representative OCT images. In Figure 3, green arrows indicate PF, red arrows indicate SF, while dark-blue arrows point to AF structures. A qualitative analysis of histology and OCT images of six samples for each age group showed a larger number of PF and SF structures in the tissue of younger mice (18-day-old, 4, and 12-week-old) than in the tissue of older ones (23-week-old). The number of PFs decreased with age, which is related to a reduction in the ovarian reservation. An increase in the number of vasculature structures in both histology and OCT images (white in histology and dark in OCT) was also observed, with a few examples indicated by yellow arrows. The number of these structures increases with age, in both histology and OCT images.

**Figure 3:**
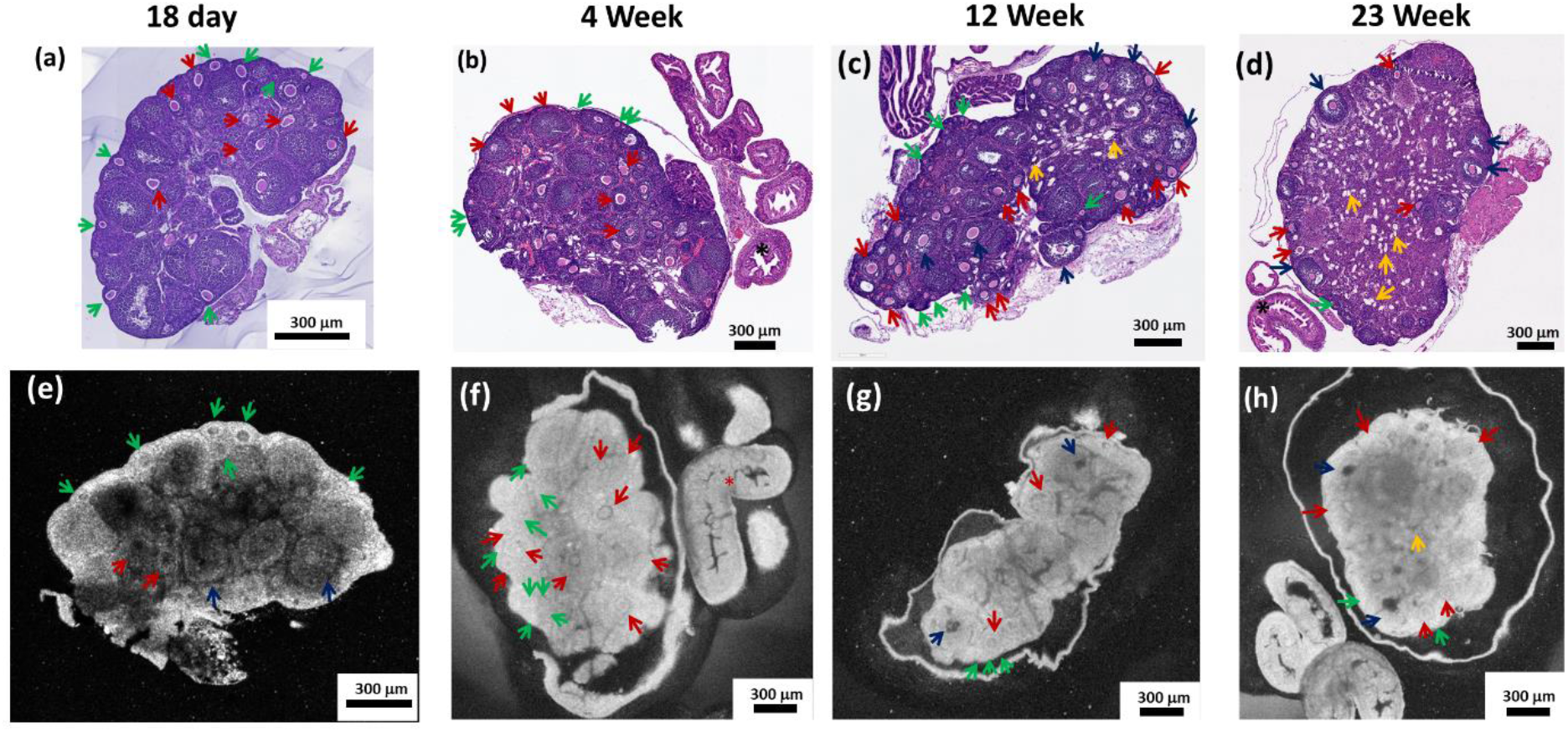
Representative H&E histology and OCT en face images of 18-day-old (a, e), 4 (b, f), 12 (c, g), and 23 (d, h) week-old mice ovarian tissue. Arrows: green, PF; red, SF; dark-blue, AF; and yellow, vessel structures.

### Attenuation Coefficient Map analysis

For comparison, Figure 4 presents OCT images before and after applying the ACM (Figure 4a and b, en face view; c and d, cross-section view). The red arrows in Figure 4 point to some of the PF. The ACM corrected the shadows in OCT images recovering the information in the affected follicles, highlighted by the rectangle en face view (Figure 4a and b), and by the asterisk mark in the cross-sectional view (Figure 4c and d). Figure 4e and f show zoomed-in views of the red rectangle in Figure 4c and d, respectively. Figure 4g and h present the A-scan signal profile obtained over the dashed line in Figure 4e and f, showing the corrected exponential decay along penetration depth after applying the ACM. Figure 4i and j show the zoomed-in views of PF’s regions (green rectangle in Figure 4c and d). The ACM images provide more structure details than conventional OCT images. Figure 4k compares the normalized signal of the granulosa cell layer of PFs at the surface and deeper image region for both OCT and ACM. The granulosa cell layer signal in OCT reduces for the deeper region while the ACM signal of the granulosa cell layer keeps stable at both surface and deeper region. The histogram of these values (Figure 4l) shows a narrower distribution for ACM (in red) normalized signal compared to OCT (gray) unveiling the decay and showing effect correction.

**Figure 4:**
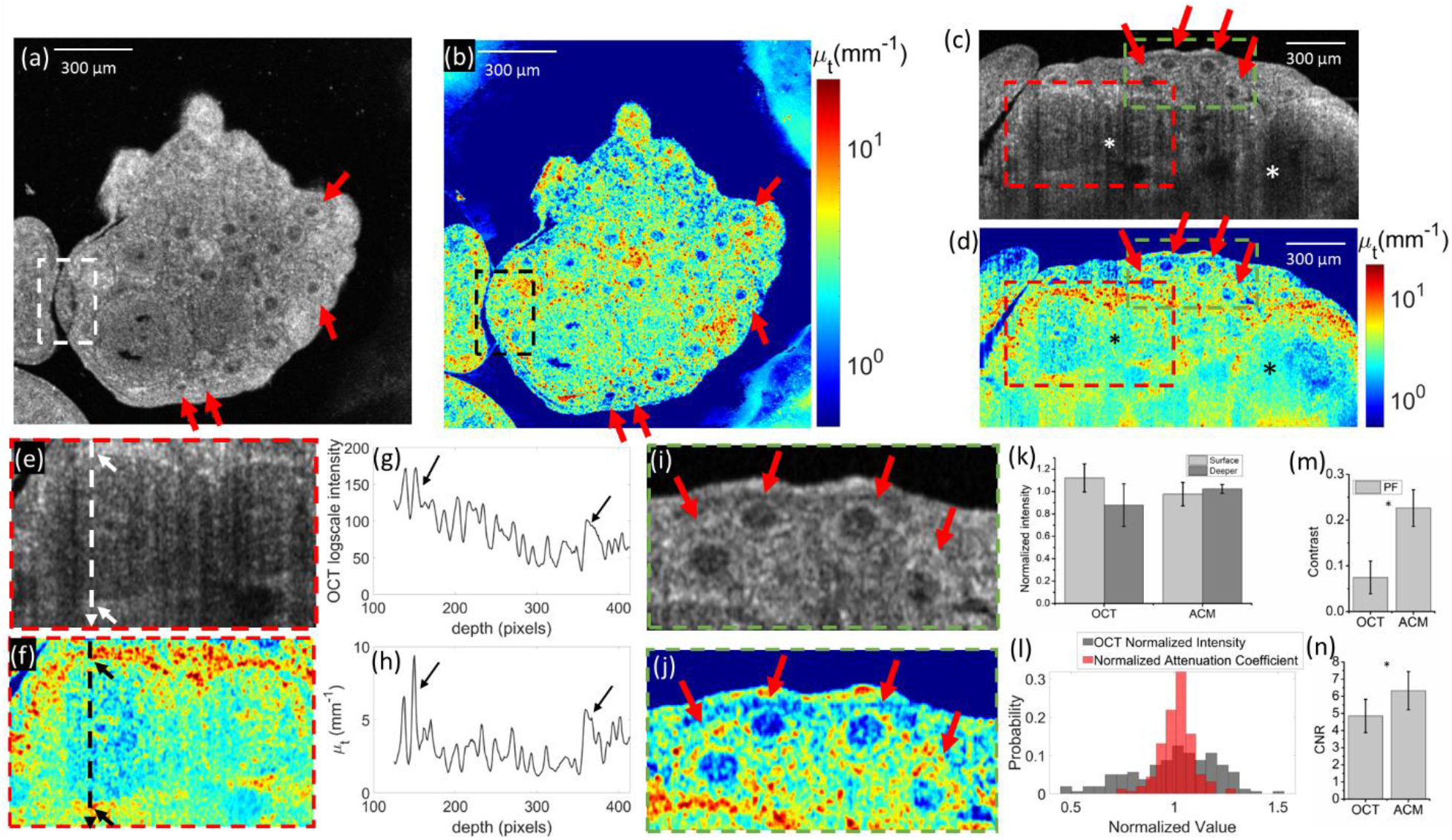
En face OCT image before (a) and after (b) applying the ACM correction. Cross-section OCT image before (c) and after (d) applying the ACM correction. (e, f) Details of Antral follicle obtained from (c,d). (g, h) Average A-scan over the dashed line position in (e, f) showing the signal correction provided by ACM. (i, j) Details of primary follicle before and after applying the ACM, obtained from (d,d) images. (k) Normalized signal of the granulosa cell layer of PFs at the surface and deeper image position. (l) Histogram of the normalized signal for OCT and ACM. (m) Image contrast and (n) contrast-to-noise ratio (CNR) analysis for primary follicle (PF) structures in OCT and ACM images.

The image improvement was also quantified by measuring the contrast (Figure 4m) and the contrast-to-noise ratio (CNR; Figure 4n) of the PF. Comparing the OCT and ACM images revealed statistically significant improvements in both the contrast (p < 0.001) and CNR (p < 0.001) of the ACM images.

### OCT imaging features that could differentiate different staged follicles

The size and the layer thickness of the granulosa cells are two features that could be potentially used to differentiate a follicle’s development stages. Figure 5a presents the diameter of each follicle stage measured in the histology and OCT images. These two measurements are in accord, confirming that the follicles were correctly identified.

**Figure 5:**
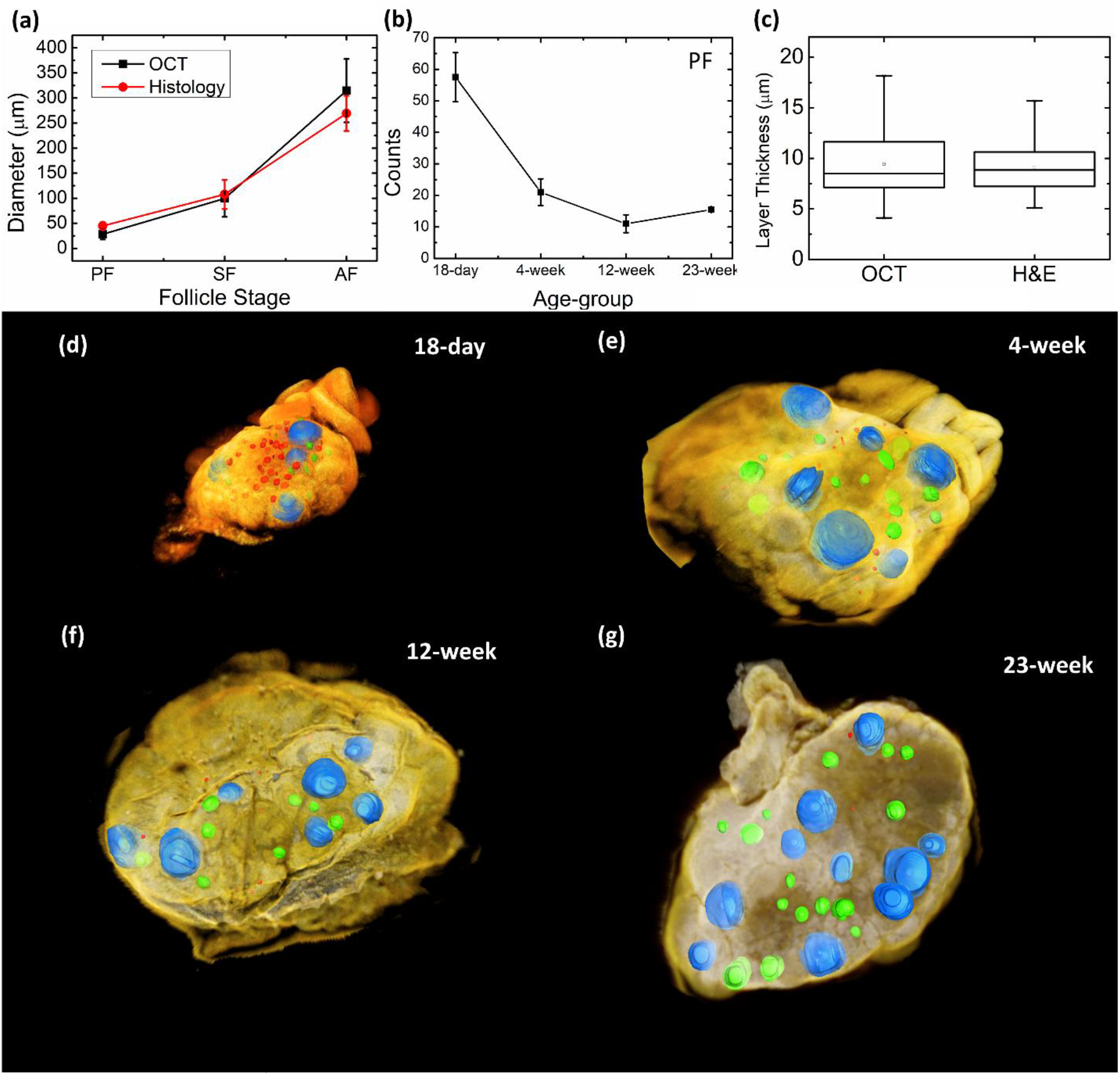
(a) Diameters of follicles in different stages, (b) follicular distribution in different age groups, and (c) follicular layers of granulosa cells thicknesses, (d, e, f, g) 3D renderings of a mouse ovary with segmented follicles in different stages (red, primary follicles; green, secondary follicles; blue, antral follicles).

The features observed in the comparison of age-related histology and OCT changes were confirmed by quantitative measurement of the number of follicles observed in the PF stage for each age group (Figure 5b). Ovarian tissues from 18-day-old mice have more PFs, reflecting their higher follicle reserve, which diminishes with age.

PFs and SFs are similar structures in the OCT image, especially in the transition from one stage to the other. The PF has one layer of granulosa cells, and SF has two or more layers of granulosa cells, which translates into different thicknesses of layers of granulosa cells in the OCT image. The layer thickness of granulosa cells is one feature that could be used to differentiate PFs from SF. Figure 5c presents the layer thicknesses of granulosa cells measured for PFs in the OCT (9.4 ± 3.8 µm) and H&E (9.1 ± 2.5 µm) images. No statistical difference in the PF layer thickness of granulosa cells was observed (p = 0.79), revealing the OCT’s ability to correctly identify this feature. OCT 3D rendering, which can capture both follicle stages, was used to segment the follicles (Figure 5d, e, f, and g) and show their locations in the ovary.

## 4. Discussion

The follicular density and stages are heterogeneously distributed over the ovarian tissue and depending on the ovarian site. Locating, classifying, and quantifying each state of the follicular development is especially important for clinical applications, such as OTC. In OTC, preserving a larger amount of follicles in their initial stages increases the chances to restore the patient fertility. Except for histological evaluation, which compromises the follicle integrity, there are no available techniques to correctly detect, identify, and quantify primordial follicles in ovarian tissue in a non-destructive fashion. A method capable of identifying and quantifying the ovarian follicles, and that does not compromise its long-term survival and grow is desirable. For this reason, OCT is a promising technique for this application.

It has been shown that OCT is a unique method for assessing ovarian tissue characterization without fixation, sectioning, and staining. In the mouse ovary, Tekae et al. demonstrated full-field OCT (FF-OCT) follicle images at each developmental stage and verified the utilities of OCT to estimate *in vitro* fertilization (IVF) outcomes in transplanted mice ovary, like ovarian reserve tests^17^. They confirmed that OCT examination did not affect IVF outcome, birth defect rate, and reproductive ability in mice. Their results also showed that the FF-OCT can correctly identify follicle developmental stages. Unfortunately, FF-OCT could not be used to accurately quantify the follicle number for the whole ovary, because the maximum imaging depth was limited to only 100 µm, with an 800 µm field-of-view (FOV). A larger effective FOV could be achieved by stitching several overlapping images together in post-processing, but this process would require a longer total data acquisition time and increase the risk of introducing image artifacts. Thus, the 3D information is limited to the superficial tissue, which restricts its clinical use. Using speckle variance OCT, Watanabe et al. enhanced the imaging contrast of oocytes in follicles, due to their microscopic movement, which could be helpful for image segmentation ^18^. In human ovarian tissue specimens, the ovary reserve was investigated using FF-OCT^19^. Due to the limited imaging depth of FF-OCT, sections of ovary cortexes were imaged, which demonstrated good morphological agreement with corresponding histology. However, this approach may have limited clinical utilities due to the limited imaging depth and lack of 3D imaging capabilities.

Therefore, in the present study, we use SD-OCT to image mouse ovaries in 3D from different age groups and identify follicles at different developmental stages. Compared with FF-OCT^17^, SD-OCT provides a much larger FOV, enabling us to image the entire mouse ovary and quantify the follicle’s sizes and density across different ages and stages.

For effective cryopreservation, it is important to correctly identify and differentiate the initial stage of development, especially the primary and secondary stages. Currently, there are no clear criteria to classify the stages of follicle maturity using OCT. The detection criteria for OCT images may depend on the visual inspection of each person operating the system, which involves a considerable amount of time and effort. Therefore, clear criteria to detect follicles according to their maturation stages from OCT images are required. Saito et al. applied a convolutional neural network^20^ to FF-OCT images to segment follicles in four-day-old mice. The authors reported a detection rate of 81 % and a precision of 67 %, but this result was restricted to two dimensions. They did not explore the 3D capability of OCT, and did not differentiate their maturity stages. In the present work, we demonstrated that OCT can be used to measure the layers of granulosa cells’ thickness of the follicle, and that this measurement can be used for correct classification.

Consistent with earlier reports, our OCT findings were well correlated with histological data, revealing follicle structures in different stages and their age-related distribution differences. We used the OCT 3D capability to measure features in the OCT image, follicle size and the layer thickness of granulosa cells, which correlated to H&E image measurements and are relevant for the correct PF classification. Moreover, our results demonstrated a strong correlation between the number of PF and age of the donor mouse, where samples from older mice had fewer PFs. Therefore, we concluded that OCT examination could be used to estimate the PF ovarian reserve and localization. This is an important capability considering future PF detection in human ovarian tissue, where the PF density is lower compared to mouse tissues. The 3D information in the OCT image was used to segment the different follicles stages for each age group (the 3D rendering is presented in the Supplementary Video).

In addition to 3D tissue structural information, OCT can provide additional contrast to enhance the identification of follicle structures. Abe et al.^18^ showed that, due to cellular activity, an oocyte can be highlighted by speckle variance OCT (SV-OCT). This approach cannot differentiate different follicular developmental stages despite their promising results. We demonstrate for the first time the use of ACM analysis to improve the contrast of mouse follicles in OCT images. This technique is independent of the signal intensity, which may vary with depth in OCT images. The ACM technique estimates the attenuation coefficient of biological tissue and enables removing shadowing artifacts from OCT images. The attenuation coefficient differences in every ovarian structure are related to different cellular density within the tissue. Compared to the original image, the obtained images have significantly higher contrast and contrast-to-noise ratio, facilitating the identification of primary follicles. Moreover, obtaining the ACM map allows us to obtain a map of intrinsic optical properties of the sample, making the method more general and independent of the OCT system setup.

## 5. Conclusions

In summary, we demonstrate the 3D imaging capability of SD-OCT to identify and quantify follicles in mouse ovaries. Follicles in three stages of development (PF, SF, and AF) were identified, and their diameters and densities were measured. The observed follicular structures and sizes match well with corresponding histological images. The layer thickness of granulosa cells was measured and used to differentiate PFs from SFs. Age-related reductions in PF follicle density was characterized using 3D OCT imaging data. Finally, we demonstrated the use of ACM analysis to improve the image contrast, facilitating the identification of primary follicles.

## Acknowledgment

This work was supported by a start-up fund from Washington University in St. Louis and NIH grants R01EB025209 and 1R21EB03268401A1. The authors thank Mr. James Ballard, and Ms. Abby Matt for thoughtful discussions and language improvement.

